# Nsp1 proteins of human coronaviruses HCoV-OC43 and SARS-CoV2 inhibit stress granule formation

**DOI:** 10.1101/2022.05.02.490272

**Authors:** Stacia M. Dolliver, Mariel Kleer, Maxwell P. Bui-Marinos, Shan Ying, Jennifer A. Corcoran, Denys A. Khaperskyy

## Abstract

Stress granules (SGs) are cytoplasmic condensates that often form as part of the cellular antiviral response. Despite the growing interest in understanding the interplay between SGs and other biological condensates and viral replication, the role of SG formation during coronavirus infection remains poorly understood. Several proteins from different coronaviruses have been shown to suppress SG formation upon overexpression, but there are only a handful of studies analyzing SG formation in coronavirus-infected cells. To better understand SG inhibition by coronaviruses, we analyzed SG formation during infection with the human common cold coronavirus OC43 (HCoV-OC43) and the highly pathogenic SARS-CoV2. We did not observe SG induction in infected cells and both viruses inhibited eukaryotic translation initiation factor 2α (eIF2α) phosphorylation and SG formation induced by exogenous stress (e.g. sodium arsenite treatment). Furthermore, in SARS-CoV2 infected cells we observed a sharp decrease in the levels of SG-nucleating protein G3BP1. Ectopic overexpression of nucleocapsid (N) and non-structural protein 1 (Nsp1) from both HCoV-OC43 and SARS-CoV-2 inhibited SG formation. The Nsp1 proteins of both viruses inhibited arsenite-induced eIF2α phosphorylation, and the Nsp1 of SARS- CoV2 alone was sufficient to cause decrease in G3BP1 levels. This phenotype was dependent on the depletion of cytoplasmic mRNA mediated by Nsp1 and associated with nuclear retention of the SG-nucleating protein TIAR. To test the role of G3BP1 in coronavirus replication, we infected cells overexpressing EGFP-tagged G3BP1 with HCoV-OC43 and observed a significant decrease in infection compared to control cells expressing EGFP. The antiviral role of G3BP1 and the existence of multiple SG suppression mechanisms that are conserved between HCoV-OC43 and SARS-CoV2 suggest that SG formation may represent an important antiviral host defense that coronaviruses target to ensure efficient replication.

**Author Summary:** Host cells possess many mechanisms that can detect viral infections and trigger defense programs to suppress viral replication and spread. One of such antiviral mechanisms is the formation of stress granules – large aggregates of RNA and proteins that sequester viral components and cellular factors needed by the virus to replicate. Because of this threat, viruses evolved specific mechanisms that prevent stress granule formation. Understanding these mechanisms can reveal potential targets for therapies that would disable viral inhibition of stress granules and render cells resistant to infection. In this study we analyzed inhibition of stress granules by two human coronaviruses: the common cold coronavirus OC43 and the pandemic SARS-CoV2. We have demonstrated that these viruses employ at least two proteins – nucleocapsid protein (N) and the non-structural protein 1 (Nsp1) to suppress stress granules. These proteins act through distinct complementary mechanisms to ensure successful virus replication. Because both OC43 and SARS-CoV2 each dedicate more than one gene product to inhibit stress granule formation, our work suggests that viral disarming of stress granule responses is central for a productive infection.

## Introduction

Coronaviruses are a family of human and animal enveloped viruses with positive-sense RNA genomes. In humans, coronaviruses predominantly cause respiratory infections of varied severity. Four circulating seasonal common cold coronaviruses (HCoVs) belong to Alphacoronavirus and Betacoronavirus genera and include HCoV-NL63, HCoV-229E, HCoV-HKU1, and HCoV-OC43 (1). In the last two decades, three novel Betacoronaviruses have entered human circulation from zoonotic sources and caused infections with high morbidity and mortality. This group includes the severe acute respiratory syndrome coronavirus (SARS-CoV), the Middle East respiratory syndrome coronavirus (MERS-CoV), and the SARS-CoV2 virus that appeared in late 2019 and caused the most devastating respiratory virus pandemic since the 1918 Spanish Flu (2–5).

The relatively genomes of Betacoronaviruses are ∼30 kb and encode 4-5 structural proteins and 16 non-structural proteins (Nsp1-16) (1,6–8). The non-structural proteins are synthesized from capped and polyadenylated genomic RNA as a polyprotein encoded by a large open reading frame (ORF), ORF1ab. This polyprotein is proteolytically processed into mature proteins by two viral enzymes: Nsp3 papain-like proteinase (PLpro) and Nsp5 3C-like proteinase (3CLpro) (1, 9). Non-structural proteins include factors that enable viral replication in the host cell and the subunits of the RNA-dependent RNA polymerase (6). Structural proteins are encoded by a nested set of subgenomic mRNAs produced by viral polymerase.

These subgenomic mRNAs also encode a variable number of smaller accessory ORFs depending on the virus species (8). Structural proteins include membrane (M), envelope (E), nucleocapsid (N), and spike (S). Of these, M and E are responsible for virus particle formation, N is an RNA-binding protein that packages viral genomes into virions, and S is the receptor binding glycoprotein that protrudes from the virion envelope and mediates membrane fusion and viral entry (10). HCoV-OC43 (OC43) S binds sialic acid that is abundant on the surface of most cell types (11), while SARS-CoV and SARS-CoV2 (CoV2) S bind angiotensin converting enzyme 2 (ACE2), determining viral tropism for the ACE2-expressing cells (12, 13).

Once virus enters a host cell, its genome associates with host translation machinery to initiate synthesis of viral proteins involved in subgenomic mRNA production, genome replication, and subversion of intrinsic host antiviral responses (14). Cytoplasmic replication of coronaviruses can generate double-stranded RNA (dsRNA) intermediates that are important pathogen-associated molecular patterns (PAMPs) sensed by the host cell (14, 15). Coronavirus dsRNA is believed to be shielded in membrane bound replication transcription compartments to limit activation of cytosolic sensors (1, 8). This mechanism, however, is insufficient to fully prevent viral dsRNA sensing, because when other mechanisms of suppression of intrinsic antiviral responses are inactivated, antiviral responses are induced in coronavirus-infected cells (16, 17). The retinoic acid inducible gene I (RIG-I) and melanoma differentiation-associated protein 5 (MDA5) can detect viral RNAs and signal to transcriptionally induce antiviral cytokines such as type I and type III interferons (15,18,19). In addition, viral replication intermediates can be recognized in cytosol by the dsRNA activated protein kinase (PKR) - one of the four kinases that can trigger inhibition of protein synthesis by phosphorylation of the α subunit of the eukaryotic translation initiation factor 2 (eIF2α) (20). When eIF2α is phosphorylated, it stably associates with the guanine exchange factor eIF2B, preventing regeneration of translation initiation-competent GTP-bound form of eIF2 (21). This blocks translation initiation and, since viruses rely on host translation for their protein synthesis, it can block viral replication. In addition, inhibition of translation initiation can induce formation of stress granules (SGs) (22–24).

SGs are large cytoplasmic condensates that accumulate translationally inactive messenger ribonucleoprotein complexes and dozens of proteins and other molecules (23, 24). SG condensation is driven by the SG-nucleating proteins like the Ras-GTPase-activating protein SH3-domain-binding protein 1 (G3BP1), G3BP2, T-cell internal antigen 1 (TIA-1) and T-cell internal antigen related (TIAR) (23–26). In addition to these and other SG-nucleating proteins, SG condensates accumulate mRNAs and translation initiation factors. In virus-infected cells, SGs may sequester viral RNAs and proteins to either disrupt their functions in the virus replication cycle or to facilitate detection of viral RNA by PAMP sensors (e.g. RIG-I) (27–31). Thus, accumulating evidence indicates that SGs are antiviral, and many viruses evolved dedicated mechanisms that inhibit their formation (32).

Several coronavirus gene products have been reported to inhibit SG formation. Of these, the most well characterized is the N protein of CoV2 that upon ectopic overexpression, directly binds G3BP1 protein and blocks SG formation induced by variety of exogenous stressors (33–35). In addition, Nsp15 protein of infectious bronchitis virus (IBV) was shown to inhibit SG formation (17). Nsp15 is a nuclease that is conserved in coronaviruses. It preferentially cleaves polyuridine RNA sequences and inhibits accumulation and detection of viral dsRNA in infected cells (36).

Another viral protein that was shown to affect SG formation is Nsp1, a host shutoff factor encoded by the first N-terminal portion of ORF1ab of betacoronaviruses (37). Nsp1 functions primarily through inhibition of host protein synthesis by binding to the 43S small ribosomal subunit complex and blocking the mRNA entry channel of the mature 80S ribosome (37–39). In addition, Nsp1 proteins of SARS-CoV and CoV2 induce host mRNA degradation by a yet to be identified host nuclease (40–42). Upon ectopic overexpression, SARS-CoV Nsp1 was shown to interact with G3BP1 and modify composition of sodium arsenite (As) induced SGs by diminishing G3BP1 recruitment (43). G3BP1 and other SG nucleating proteins drive SG formation by interacting with polysome-free messenger ribonucleoproteins that accumulate after inhibition of translation initiation. Therefore, in theory, inhibition of translation initiation by Nsp1 could promote SG formation by causing polysome disassembly. At the same time, host mRNA degradation could deplete untranslated mRNAs and inhibit SG formation, and we have shown previously that the influenza A virus host shutoff endonuclease PA-X can potently inhibit SG formation through this mechanism (31). Which of the two Nsp1 host shutoff features is more dominant in modulating SGs and how Nsp1 proteins that do not induce mRNA degradation affect SG formation has not been tested to date.

Given the proposed antiviral functions of SG formation and the multiple proposed mechanisms that betacoronaviruses employ to block SG formation and modify their composition, we aimed at characterizing SG inhibition mechanisms by the model common cold betacoronavirus OC43. We compared SG inhibition by this virus and the highly pathogenic CoV2 in the same cell culture infection model to identify potential common strategies and/or differences in the magnitude and molecular mechanisms of SG inhibition. In this work we demonstrate that both OC43 and CoV2 efficiently inhibit SG formation in infected cells and show that N and Nsp1 proteins of both viruses act through distinct mechanisms to inhibit SG formation. N proteins of OC43 and CoV2 act independent from eIF2α phosphorylation and downstream of translation arrest, while Nsp1 proteins block SG formation by inhibiting eIF2α phosphorylation upstream of SG nucleation. In addition, CoV2 but not OC43 infection causes depletion of G3BP1 and disrupts nucleocytoplasmic shuttling of TIAR, which contributes to more potent inhibition of SG formation by this virus. We demonstrate that the CoV2 Nsp1-mediated host shutoff is responsible, at least in part, for depletion of G3BP1 and nuclear accumulation of TIAR. Specifically, the mRNA degradation function was required for these phenotypes. When we overexpressed G3BP1, it significantly decreased OC43 infection, illustrating that G3BP1 is antiviral towards coronaviruses. Our study reveals that both OC43 and CoV2 each dedicate more than one gene product to inhibit SG formation, a genetic redundancy that supports that viral disarming of SG responses is central for a productive infection.

## Results

### Human coronavirus OC43 inhibits SG formation in infected cells

Previously in our laboratory we established a robust OC43 infection model in human embryonic kidney (HEK) 293A cells (44). This cell line, historically used for isolation and titration of adenoviruses (45–47), is also readily infected with many RNA viruses, including OC43, which rapidly replicates in 293A cells to high titers. To examine SG dynamics over the course of infection, we infected 293A cells with OC43 at multiplicity of infection (MOI) of 1.0 and analysed infected cells for the presence of SGs at various times post-infection using immunofluorescence staining for TIAR protein (Fig. 1A). We observed little to no SG formation until 48 h post infection (hpi), by which time many cells started lifting off and dying from infection. On average, only 5% of infected cells formed SGs at 24 hpi (Fig.1A,D). Because there was little SG formation in OC43-infected cells, we analysed if OC43 actively inhibited SG formation. We treated mock and virus-infected cells with sodium arsenite (As) at 24 hpi. Arsenite is a potent SG-inducing agent which is commonly used to induce high levels of eIF2α phosphorylation and SG formation. It causes oxidative stress and activates eIF2α kinase heme-regulated inhibitor (HRI) (22, 25). As expected, in mock-infected cells, SGs were induced in nearly 100% of cells. By contrast, less than half of the OC43-infected cells formed SGs following As treatment (Fig.1B,D). Next, we tested levels of eIF2α phosphorylation in infected cells and discovered that OC43 infection inhibited As-induced eIF2α phosphorylation, with increasing efficiency from 12 to 48 hpi (Fig.1C). Thus, the inhibition of SG formation in OC43 infected cells could be, at least in part, due to viral inhibition of eIF2α phosphorylation-induced translation arrest upstream of SG nucleation.

**Figure 1.**
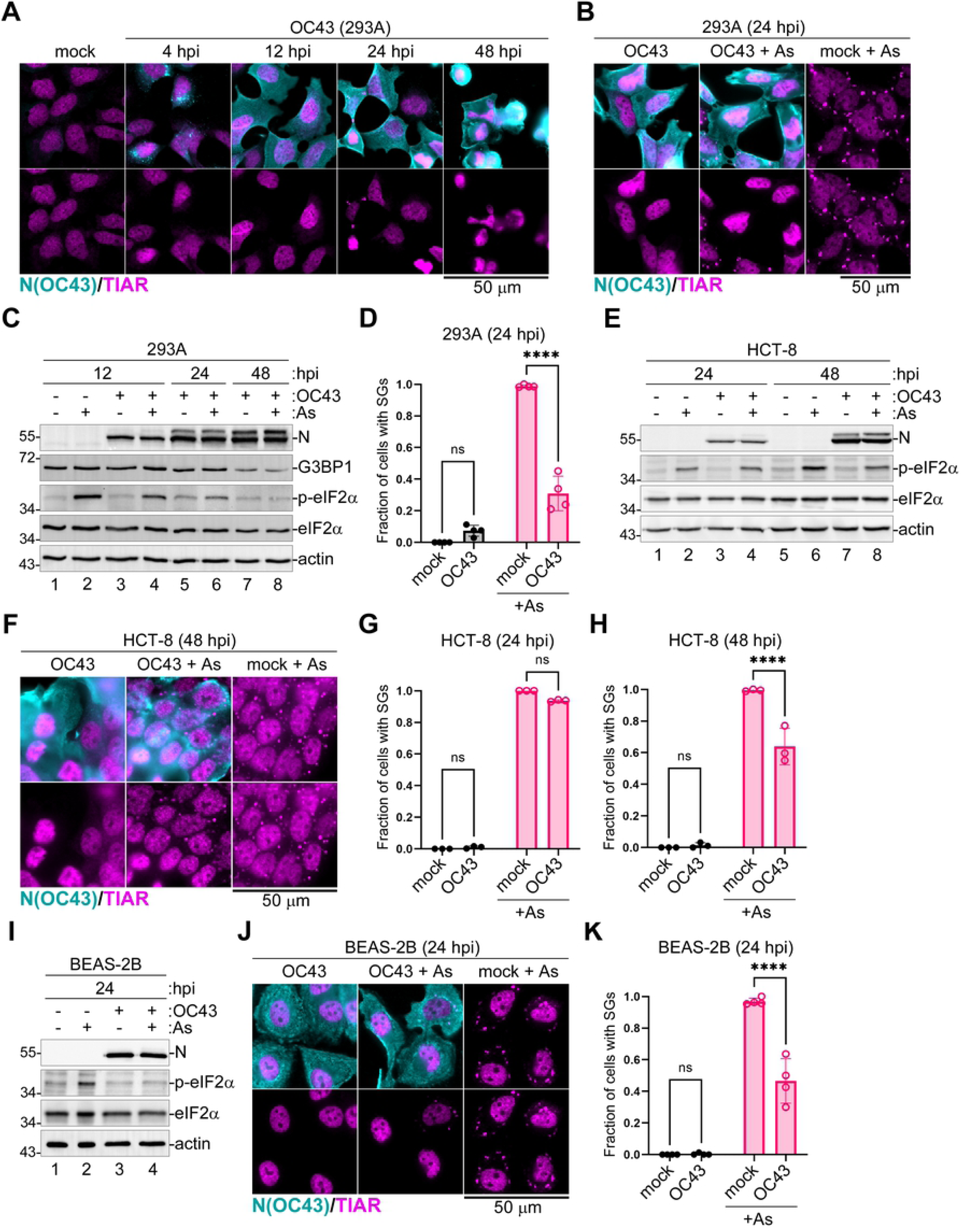
Coronavirus OC43 inhibits SG formation and eIF2α phosphorylation. Cells were infected with OC43 and SG formation in infected cells was analyzed at the indicated times post-infection using immunofluorescence staining for nucleoprotein (N(OC43), teal) and SG marker TIAR (magenta). Levels of N protein accumulation and phosphorylation of eIF2α were analysed by western blot. hpi = hours post-infection. Scale bars = 50 µm. (A) Immunofluorescence analysis of infected 293A cells at different times post-infection. (B,F,J) Immunofluorescence analysis of SG formation in mock infected and OC43- infected 293A (B), HCT-8 (F), and BEAS-2B (J) cells treated with sodium arsenite (+ As) or untreated infected cells at indicated times post-infection. (C,E,I) Western blot analysis of As-induced eIF2α phosphorylation and accumulation of N protein in 293A (C), HCT-8 (E), and BEAS-2B (I) cells at the indicated times post-infection. Levels of SG nucleating protein G3BP1 were also analyzed in (C). Actin was used as loading control. (D,G,H,K) Fraction of cells with SGs was quantified in mock and OC43- infected 293A cells at the indicated times post-infection. Two-way ANOVA and Tukey multiple comparisons tests were done to determine statistical significance (****, p -value < 0.0001, ns = non-significant). On all plots each data point represents independent biological replicate (N ≥ 3). Error bars = standard deviation.

To verify that our observations are not specific for 293A cells, we repeated analyses of OC43 effects on As-induced eIF2α phosphorylation and SG formation in human colon (HCT-8) cells, which are often used to grow this virus, and the immortalized primary human upper airway epithelial BEAS-2B cells that more closely represent a cell type infected by coronaviruses *in vivo*. Arsenite-induced eIF2α phosphorylation and SG formation were inhibited in HCT-8 cells, however at the later time point, 48 hpi (Fig.1E-H), possibly reflecting slower virus replication kinetics in this cell type compared to 293A. Indeed, at 24 hpi, much lower levels of OC43 N protein were detected in HCT-8 cells (compare lanes 3 and 4 to 7 and 8 in Fig. 1E). Similar to 293A, in infected BEAS-2B cells, eIF2α phosphorylation and SG formation were inhibited at 24 hpi (Fig. 1I-K), indicating that these phenotypes are not limited to fully transformed cell types and that our 293A infection model is appropriate for analysis of SG responses to infection.

### Inhibition of SG formation in OC43-infected cells does not depend on blocking eIF2α phosphorylation

Given that OC43 infection simultaneously decreased arsenite-induced SG formation and eIF2α phosphorylation, we decided to test if this virus could block SGs formation induced by an eIF2α phosphorylation-independent pathway. To induce SGs in these experiments, we used Silvestrol, which inhibits the helicase eIF4A, an important translation initiation factor, and triggers SG formation without inducing phosphorylation of eIF2α (48, 49). 293A cells were infected with OC43 and treated with Silvestrol for 1 hour prior to analysis at 24 h post-infection. Similar to arsenite, Silvestrol treatment triggered SG formation in nearly 100% of mock-infected cells, while only half of OC43-infected cells had SGs (Fig. 2A,C). As expected, Silvestrol treatment did not induce eIF2α phosphorylation (Fig.2B).

**Figure 2.**
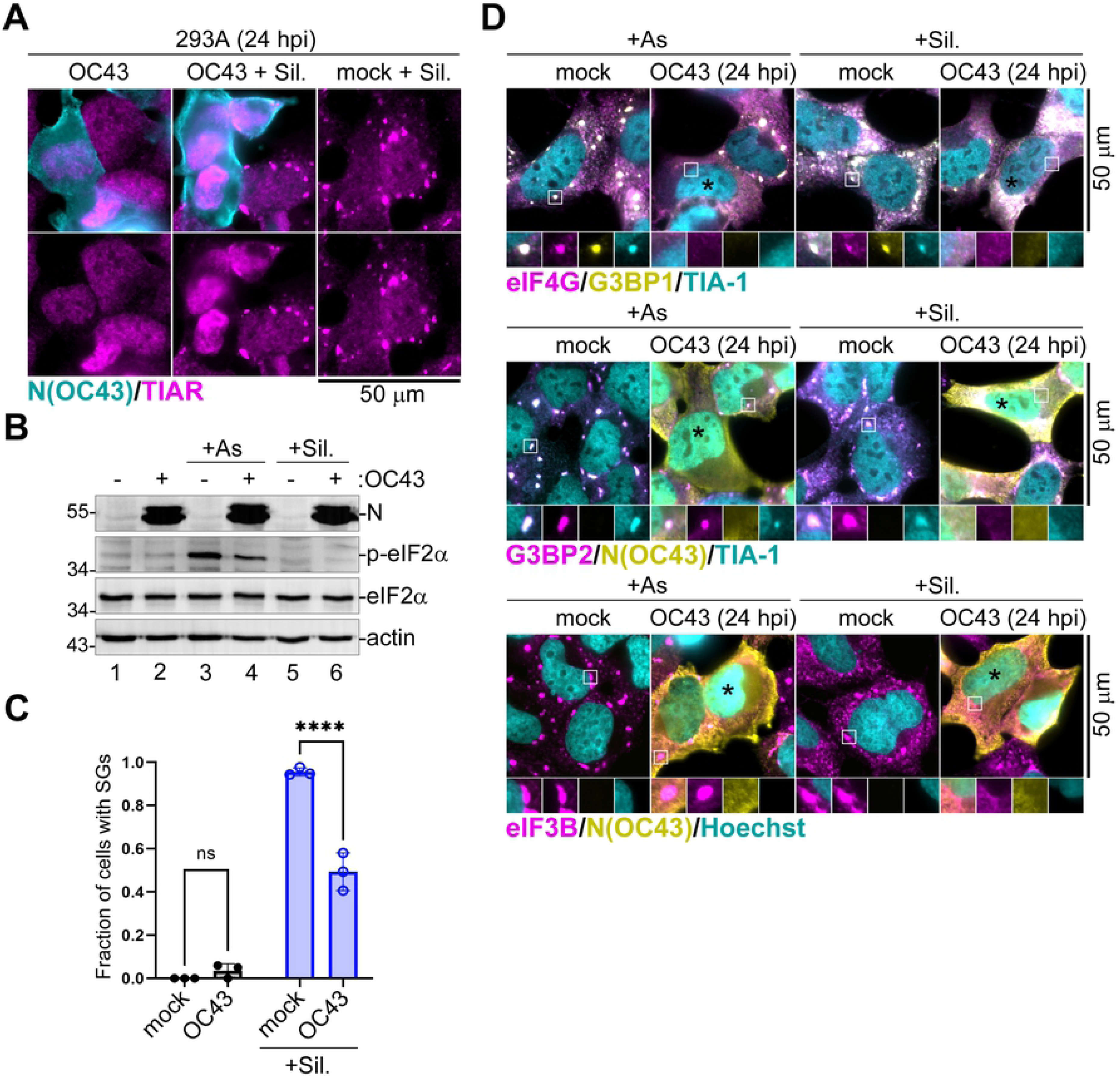
OC43 inhibits SGs independently of eIF2α phosphorylation. 293A cells were infected with OC43 at MOI = 1.0 and SG formation in arsenite (+ As), Silvestrol (+ Sil.), and untreated mock and OC43 infected cells was analyzed at 24 hours post-infection (hpi) using immunofluorescence staining for nucleoprotein and the indicated SG markers. (A) Immunofluorescence analysis of SG formation in mock infected and OC43-infected cells treated with Silvestrol (+ Sil.) or untreated infected cells. (B) Phosphorylation of eIF2α was analysed by western blot. (C) Fraction of cells with SGs was quantified in mock and OC43-infected 293A cells treated and stained as in panel (A). Each data point represents independent biological replicate (N=3). Error bars = standard deviation. Two-way ANOVA and Tukey multiple comparisons tests were done to determine statistical significance (****, p -value < 0.0001, ns = non-significant). (D) Representative images of mock infected and OC43 infected cells immunostained for SG markers eIF4G (magenta), G3BP1 (yellow), TIA-1 (teal), G3BP2 (magenta), and eIF3B (magenta) as indicated. Subcellular distribution of nucleoprotein (N(OC43), yellow) was visualised by immunostaining and nuclear DNA was visualised with Hoechst dye (teal) where indicated. Black asterisks indicate infected cells that did not form SGs. Outsets show enlarged areas of cytoplasm with separation of channels to better visualize co-localization of SGs markers. Scale bars = 50 μm.

Interestingly, in many infected cells we noticed brighter nuclear TIAR staining compared to uninfected cells, possibly indicating disruption of normal nucleocytoplasmic shuttling of TIAR (Fig. 1A,B, Fig. 2A). To confirm that OC43 effects were not limited to TIAR-containing SGs, we completed a series of experiments using multiple SG marker proteins to analyze SG formation in infected cells treated with either As or Silvestrol. These analyses confirmed that regardless of the markers used, including SG-nucleating proteins G3BP1, G3BP2, and TIA-1, as well as translation initiation factors eIF4G and eIF3B, formation of SG foci was inhibited by OC43 (Fig.2D). In addition, these analyses revealed that in a fraction of infected cells that did form SGs, OC43 N protein was not accumulating in these foci (Fig. 2D).

### SARS-CoV2 blocks SG formation in infected cells and depletes SG nucleating protein G3BP1

Ectopic overexpression studies suggest that several gene products of pandemic CoV-2 virus can suppresses SG formation (17,35,50,51). To test if CoV2 can effectively block SG formation in our cell culture system, we infected 293A cells stably expressing ACE2 (293A-ACE2) with this virus and analysed SG formation. In our *in vitro* infection model, no SG formation was observed in CoV2-infected cells at either 12 hpi or 24 hpi (Fig.3A). In addition, CoV2 was able to effectively suppress SG formation induced by As (Fig. 3B,C). Compared to OC43, CoV2 infection resulted in much greater SG inhibition, with only about 8% of infected cells forming SGs following As treatment (Fig. 3C). Similar to OC43, CoV2 inhibited As-induced eIF2α phosphorylation (Fig. 3D). Interestingly, we also observed an increase in nuclear TIAR signal in CoV2 infected cells, which was even more prominent than that was observed in OC43 infected cells (Fig.3A,B). When we compared total levels of TIAR in mock and CoV2 infected cells using western blot, we observed similar levels, indicating that the virus causes changes in subcellular distribution of TIAR without drastically affecting its expression (Fig. 3D,E). By contrast, we consistently observed substantial reduction in the levels of G3BP1 protein in CoV2 infected cells compared to mock (Fig. 3D,E). Unlike OC43 infection, which resulted in detectable decrease in G3BP1 levels only at 48 hpi (Fig. 1C), in CoV2 infected cells G3BP1 decrease was observed much earlier (Fig. 3D,E). To examine if these changes in G3BP1 expression were due to a decrease in its transcript levels, we isolated total RNA from mock and CoV2-infected cells at 24 hpi and analysed G3BP1 and TIAR mRNA levels using RT-QPCR. Consistent with a known feature of CoV2 host shutoff causing cytoplasmic mRNA degradation, we detected dramatic depletion of both G3BP1 and TIAR transcripts in infected cells (Fig. 3F). The magnitude of mRNA depletion was slightly higher for G3BP1 (on average 80% depleted for G3BP1 vs. 60% for TIAR), but alone it would not account for the observed differences in protein levels in infected cells. To examine relative stability of G3BP1 and TIAR proteins in 293A cells, we treated uninfected cells with two translation inhibitors, cycloheximide (CHX) and Silvestrol (Sil.), as well as the transcription inhibitor Actinomycin D (ActD) for 12 hours and measured protein levels using western blot. As expected, within 1 hour of treatment, CHX and Sil. potently blocked protein synthesis in 293A cells, while ActD had no effect (Fig. 3G). However, after 12 h only ActD treatment resulted in a small (∼25%) but statistically significant decrease in G3BP1 and, to a lesser extent, TIAR protein levels, while translation inhibitors did not decrease either of these proteins (Fig. 3H-J). This suggests that neither G3BP1 nor TIAR are intrinsically unstable and rapidly degraded proteins. Instead, it points to a distinct turnover mechanism for these RNA-binding proteins that is activated in response to either decrease in transcription or total mRNA levels. As a control, we examined total levels of the translation initiation factor eIF4G and detected no changes in its expression after ActD treatment (Fig. 3K). These results suggest that the dramatic decrease in G3BP1 levels in CoV2 infected cells is primarily due to mRNA depletion rather than translation arrest, which are both features of CoV2 host shutoff. However, a direct targeting of G3BP1 for degradation by a viral protein cannot be ruled out. In either case, given the central role of G3BP1 in nucleating SGs, its depletion undoubtedly contributes to potent SG suppression by CoV2.

**Figure 3.**
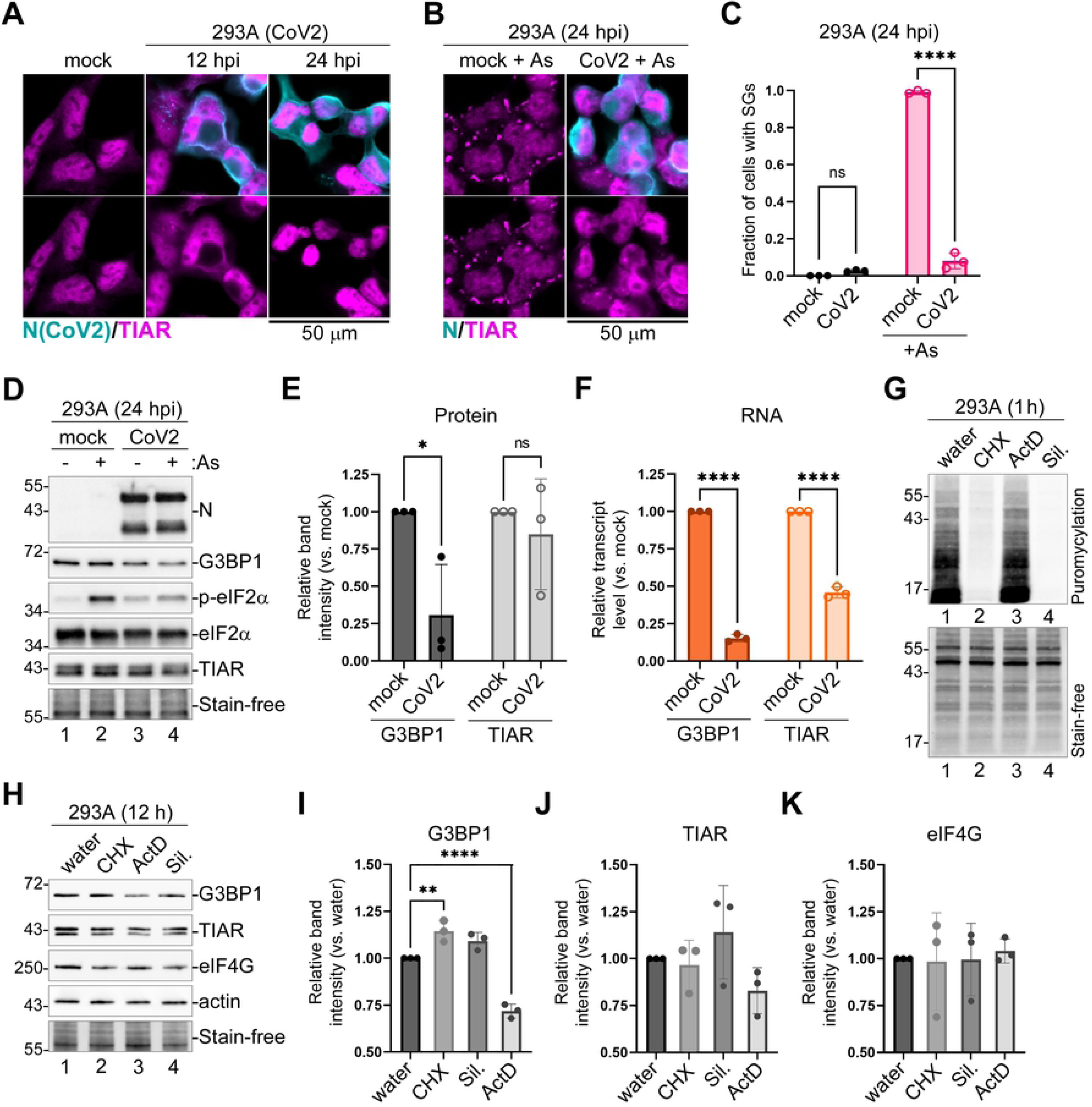
SARS-CoV-2 inhibits SG formation and decreases expression of G3BP1. (A) 293A-ACE2 cells were infected with CoV2 at MOI = 1.0 and mock and CoV2 infected cells were analysed by immunofluorescence microscopy using immunostaining for viral nucleoprotein (N, teal) and SG marker TIAR (magenta) at the indicated times post-infection. (B) Immunofluorescence microscopy of mock and CoV2 infected 293A-ACE2 cells treated with As at 24 hpi. Staining was performed as in (A). In (A) and (B) scale bars = 50 µm. (C) Fraction of cells with SGs was quantified in mock and CoV2 infected cells treated and stained as in panel (B). Error bars = standard deviation. Two-way ANOVA and Tukey multiple comparisons tests were done to determine statistical significance (****, p -value < 0.0001, ns = non-significant). (D) Western blot analysis of As-induced eIF2α phosphorylation and accumulation of N protein in 293A-ACE2 cells at 24 hpi. Levels of SG nucleating proteins G3BP1 and TIAR were also analyzed, fluorescent total protein stain (Stain-free) was used as loading control. (E) Band intensity of G3BP1 and TIAR normalized to total protein (Stain-free) quantified from 3 independent experiments represented in D (lanes 1 and 3), (*, p -value < 0.05, ns, non-significant, as determined by unpaired Students t-Test). (F) Relative transcript levels of G3BP1 and TIAR determined by RT-QPCR (****, p - value < 0.0001, ns, non-significant, as determined by unpaired Students t-Test). (G) Ribopuromycylation assay in 293A cells treated with cycloheximide (CHX), Actinomycin D (ActD), Silvestrol (Sil.) or water control. Top - puromycin incorporation detected by western blot. Bottom – total protein visualized by Stain-free dye. (H) Western blot of 293A cells treated as in G for 12h. (I-K) Relative band intensity of G3BP1 (I), TIAR (J), and eIF4G (K) normalized to total protein (Stain-free), quantified from 3 independent experiments represented in H. Two-way ANOVA and Tukey multiple comparisons tests were done to determine statistical significance (**, p -value < 0.01; ****, p -value < 0.0001). On all plots each data point represents independent biological replicate (N=3). Error bars = standard deviation.

### N proteins of OC43 and CoV2 inhibit SG formation

Previous studies demonstrated that CoV2 N and Nsp15 proteins inhibit SG formation upon ectopic overexpression (17,33,51–53). To test if OC43 N and Nsp15 proteins contribute to SG inhibition by this virus, we overexpressed EGFP-tagged OC43 N and Nsp15 proteins, the EGFP-tagged CoV2 N protein, or EGFP control in 293A cells and assessed their effect on SG formation. First, we analyzed SG formation in cells transiently transfected with CoV2 and OC43 N expression constructs or EGFP control and treated with As at 24 h post-transfection. Western blot analysis revealed that the OC43 and CoV2 EGFP-N fusion proteins were expressed at similar levels, and that neither of the N constructs affected G3BP1 expression levels or As-induced eIF2α phosphorylation (Fig. 4A). As expected, immunofluorescence microscopy analyses showed that nearly 100% of the EGFP expressing cells formed SGs upon As treatment, while CoV2 N expression efficiently inhibited SG formation (Fig. 4B,C). In OC43 N expressing cells, SG formation was also inhibited, but to a lesser extent (Fig. 4C). Given that N proteins did not affect eIF2α phosphorylation, we next tested if OC43 and/or CoV2 EGFP-N constructs could inhibit Silvestrol-induced SG formation. Indeed, both N constructs were able to inhibit Silvestrol-induced SG formation (Fig 4D,E), and the OC43 N protein was even better at inhibiting Silvestrol-induced SGs than the As- induced SGs (Fig. 4C,E). This indicates that N proteins of these coronaviruses directly affect SG formation downstream of translation arrest, consistent with direct interaction with SG nucleating protein G3BP1 by CoV2 N reported by previous studies (33, 51).

**Figure 4.**
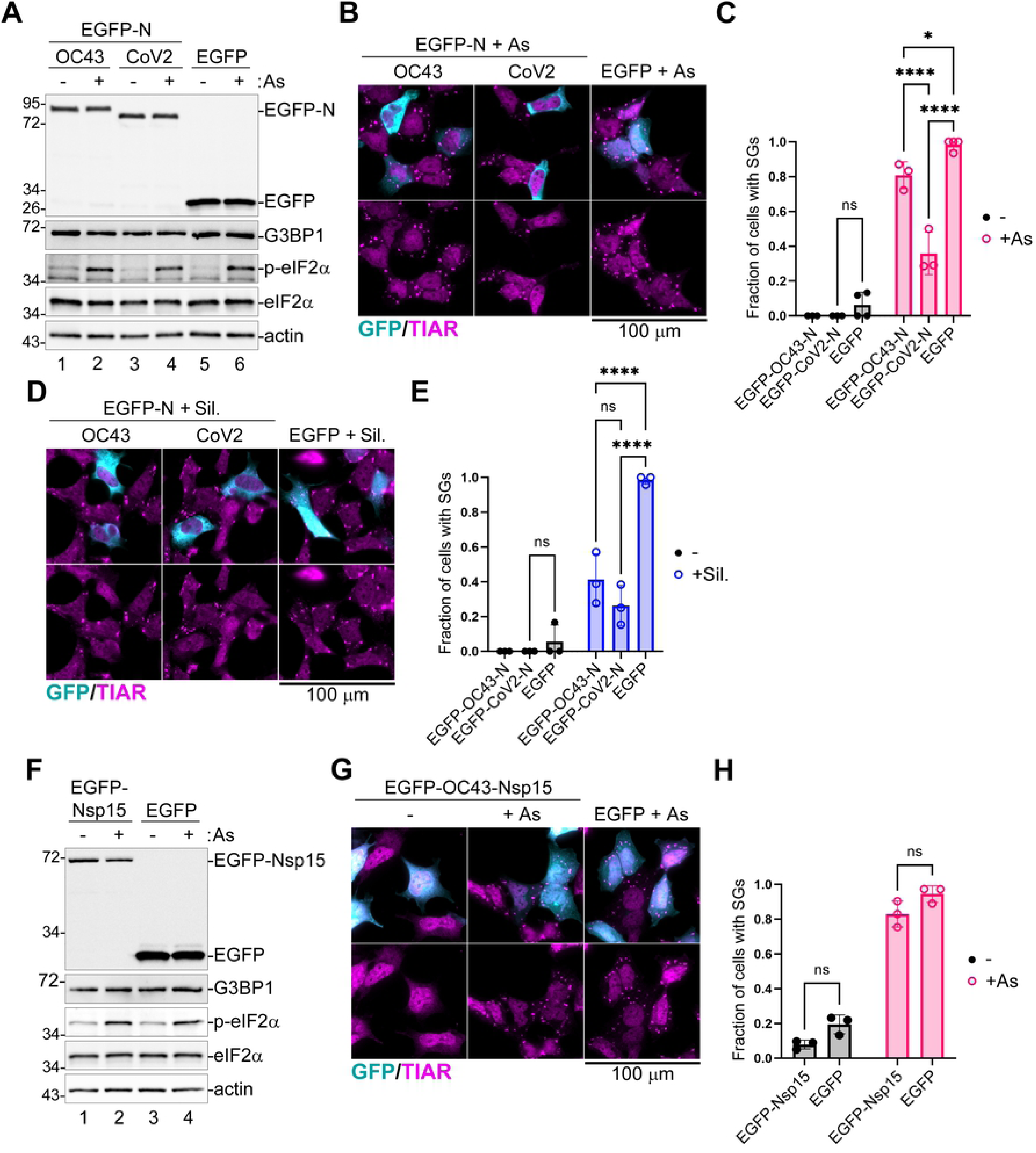
Coronavirus N proteins inhibit SG formation downstream of eIF2α phosphorylation. 293A cells were transiently transfected with the indicated EGFP-tagged viral protein expression constructs or EGFP control. At 24 h post-transfection cells were treated with sodium arsenite (+ As) or Silvestrol (+ Sil.), as indicated, or left untreated. SG formation was analyzed by immunofluorescence microscopy with staining for SG marker TIAR (magenta). EGFP expression is shown in teal. Scale bars = 100 μm. Levels of EGFP-tagged proteins and the As-induced phosphorylation of eIF2α were analyzed by western blot. (A) Western blot analysis of cells transfected with EGFP-tagged OC43 or CoV2 nucleoprotein (EGFP-N) expression vectors or control EGFP-transfected cells. (B) Immunofluorescence microscopy of EGFP-N transfected, or control EGFP-transfected cells treated with arsenite. C) Fraction of transfected cells with As-induced SGs quantified from B. (D) Immunofluorescence microscopy of EGFP-N transfected or control EGFP-transfected cells treated with Silvestrol. (E) Fraction of transfected cells with Silvestrol-induced SGs quantified from D. (F) Western blot of OC43 EGFP-Nsp15 transfected or control EGFP transfected cells treated with arsenite. (G) Immunofluorescence microscopy of OC43 EGFP-Nsp15 transfected, or control EGFP-transfected cells treated with arsenite. (H) Fraction of transfected cells with As-induced SGs quantified from G. On all plots each data point represents independent biological replicate (N=3). Error bars = standard deviation. Two-way ANOVA and Tukey multiple comparisons tests were done to determine statistical significance (****, p -value < 0.0001, *, p- value < 0.05, ns, non-significant).

Next, we focused on Nsp15 of OC43 to determine if it can interfere with SG formation. We transfected 293A cells with EGFP-tagged OC43 Nsp15 or EGFP control and treated cells with As at 24 h post- transfection to induce SGs. Similar to N constructs, EGFP-Nsp15 did not affect As-induced eIF2α phosphorylation (Fig. 4F). Furthermore, OC43 Nsp15 did not significantly affect As-induced SG formation (Fig. 4G,H). Thus, we conclude that Nsp15 is not contributing to inhibition of As-induced SGs by OC43 in our experimental system.

### Nsp1-mediated host shutoff contributes to SG inhibition by coronaviruses

N proteins of both OC43 and CoV2 coronaviruses are responsible, at least in part, for inhibition of SG condensation downstream of translation arrest, without affecting levels of eIF2α phosphorylation or expression of G3BP1. However, the magnitude of SG inhibition observed in EGFP-N expressing cells was much lower than in virus-infected cells. Therefore, it is likely that additional mechanisms of SG suppression may be used by these coronaviruses. Because of the link between host mRNA degradation mediated by CoV2 host shutoff and the depletion of G3BP1, we tested if Nsp1 proteins, major host shutoff factors of OC43 and CoV2, could inhibit SG formation. Importantly, only CoV2 Nsp1 was previously shown to induce degradation of cellular mRNAs (39,42,54), while both OC43 and CoV2 Nsp1 proteins block translation initiation. When we transfected 293A cells with N-terminally HA-tagged Nsp1 constructs or EGFP control and induced SG formation with As 24 h post-transfection, we saw that both OC43 and CoV2 Nsp1 were inhibiting SG formation as determined by TIAR staining (Fig. 5A). We also observed that CoV2 Nsp1 was causing depletion of cytoplasmic and increase in nuclear TIAR signal. This suggests that Nsp1 is responsible for disruption of nucleocytoplasmic shuttling of TIAR we saw previously in CoV2 infected cells. Western blotting analysis demonstrated that our Nsp1 constructs were inhibiting production of co-transfected EGFP reporter, as expected, confirming their activity in blocking host gene expression. Notably, CoV2 Nsp1 had stronger effect on EGFP expression, likely because of additional activity of stimulating mRNA degradation (Fig. 5B). In addition, western blot analysis revealed that both OC43 and CoV2 Nsp1 significantly attenuated As-induced eIF2α phosphorylation (Fig. 5B,C). OC43 Nsp1 inhibited eIF2α phosphorylation by 30% on average, and CoV2 Nsp1 by 40%. Since we consistently observed transfection efficiencies of 40-60% in these experiments, inhibition of p-eIF2α in transfected cells may be even stronger. Next, we tested if Nsp1 proteins affected G3BP1 protein expression. Because our HA and G3BP1 antibodies suitable for immunofluorescence staining are both mouse, we constructed N-terminally EGFP-tagged Nsp1 constructs and analysed levels and subcellular localization of G3BP1 in transfected cells treated with As. In OC43 EGFP-Nsp1 expressing cells, SG formation was inhibited and G3BP1 was diffusely distributed in the cytoplasm even upon treatment with As. In contrast, in CoV2 EGFP-Nsp1 expressing cells, G3BP1 signal was much lower than in bystander untransfected cells, EGFP expressing control cells, or OC43 EGFP-Nsp1 cells (Fig. 5D). Both TIAR and G3BP1 are important SG nucleating proteins, therefore the CoV2 Nsp1 mediated redistribution of TIAR into the nucleus and depletion of G3BP1 levels could potentially disrupt cytoplasmic SG condensation.

**Figure 5.**
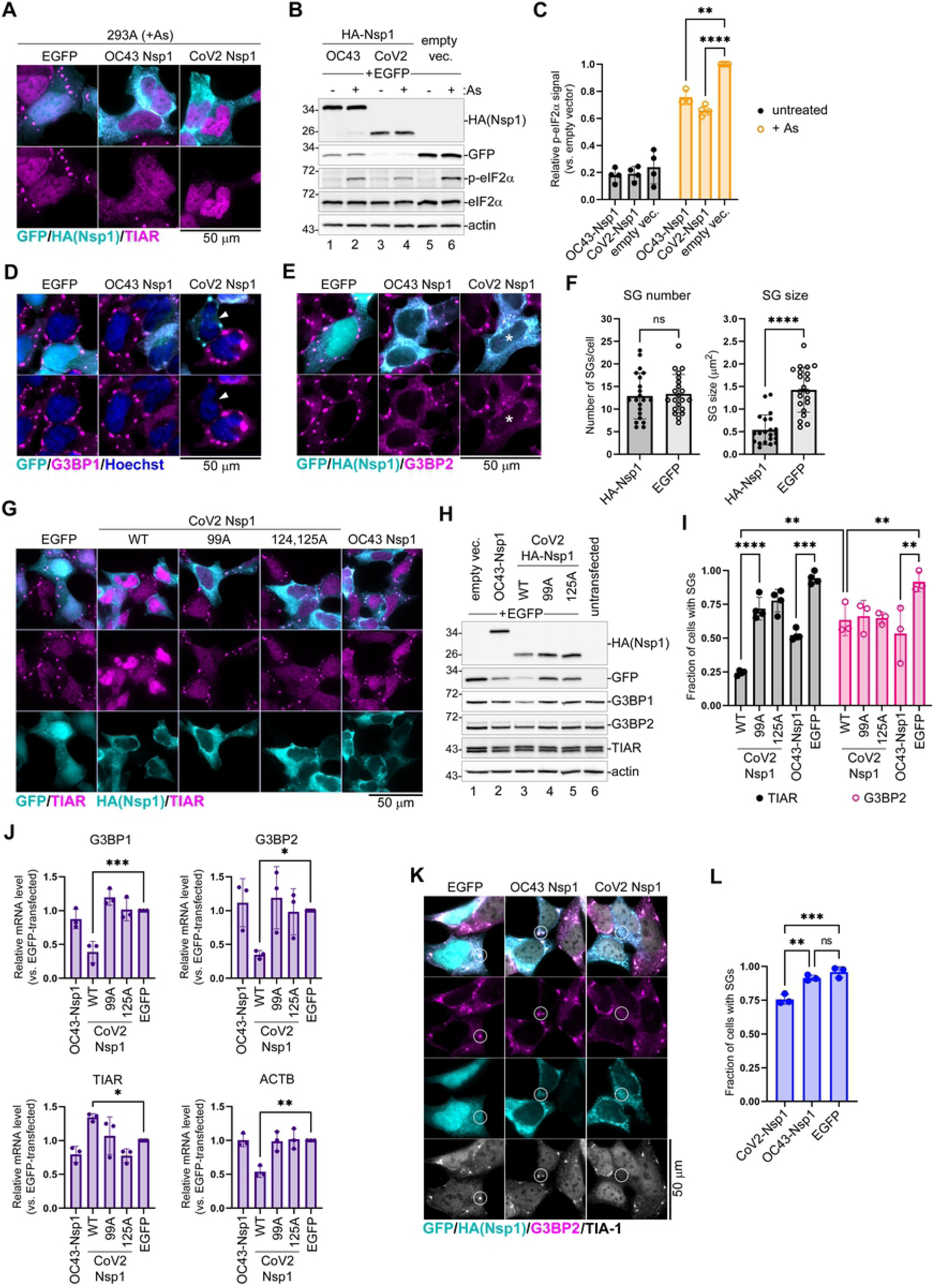
Nsp1 inhibits eIF2α phosphorylation and SG formation. (A) 293A cells were transiently transfected with the indicated HA-tagged Nsp1 expression constructs or EGFP control. At 24 h post- transfection cells were treated with sodium arsenite (+ As) and SG formation analysed by immunofluorescence microscopy staining for TIAR (magenta). GFP signal in control cells and HA- tagged Nsp1 signal are shown in teal. Scale bar = 50 μm. (B) Whole cell lysates from cells treated as in A with or without As were analysed by western blot. (C) Phospho-eIF2α band intensity normalized to total eIF2α was quantified from B and plotted relative to phospho-eIF2α level in EGFP control cells treated with As. Each data point represents independent biological replicate (N≥3). Error bars = standard deviation. Two-way ANOVA and Tukey multiple comparisons tests were done to determine statistical significance (****, p -value < 0.0001; ***, p-value <0.001; **, p-value < 0.01). (D) Immunofluorescence microscopy analysis of cells expressing the indicated EGFP-tagged Nsp1 constructs or EGFP control. SG formation was visualized by staining for G3BP1 (magenta). GFP signal is shown in teal. Arrowhead indicates a representative cell with low G3BP1 signal. Scale bar = 50 μm. (E) Immunofluorescence microscopy analysis of cells expressing the indicated HA-tagged Nsp1 constructs or EGFP control. SG formation was visualized by staining for G3BP2 (magenta). GFP signal in control cells and HA-tagged Nsp1 signal are shown in teal. Asterisks indicates a representative CoV2 Nsp1 expressing cell with small SGs. Scale bar = 50 μm. (F) G3BP2-positive SG number per cell and average SG size per cell were quantified in CoV2 HA-Nsp1-transfected cells and control EGFP-transfected cells from E. Each data point represents individual cell analysed from 3 independent biological replicates (21 cells per condition). Error bars = standard deviation. Two-tailed Students t-Test was done to determine statistical significance. (****, p -value < 0.0001, ns, non-significant). (G) Immunofluorescence microscopy analysis of cells expressing the indicated wild type and mutant Nsp1 constructs. SG formation was visualized by staining for TIAR (magenta). GFP signal in control cells and HA-tagged Nsp1 signal is shown in teal. WT = wild type; 99A = R99A mutant; 125A = R124A,K125A mutant. Scale bar = 50 μm. (H) Western blot analysis of cells co-transfected with the indicated HA-tagged Nsp1 constructs and EGFP. (I) Fraction of transfected cells with As-induced SGs quantified from G (TIAR) and E (G3BP2). Each data point represents independent biological replicate (N≥3). Error bars = standard deviation. Two-way ANOVA and Tukey multiple comparisons tests were done to determine statistical significance (****, p -value < 0.0001; **, p-value <0.01; *, p-value < 0.05, ns, non-significant). (J) Relative G3BP1, G3BP2, TIAR, and ACTB mRNA levels were determined using RT-QPCR in cells transfected with the indicated expression constructs at 24 h post-transfections. Values were normalized to 18S. Each data point represents independent biological replicate (N=3). Error bars = standard deviation. One-way ANOVA and Dunnett’s multiple comparisons tests were done to determine statistical significance (***, p -value < 0.001; **, p- value <0.01; *, p-value < 0.05). (K) Immunofluorescence microscopy analysis of cells expressing the indicated HA-tagged Nsp1 constructs or EGFP control and treated with Silvestrol. SG formation was visualized by staining for G3BP2 (magenta) and TIA-1 (greyscale). GFP signal in control cells and HA- tagged Nsp1 signal are shown in teal. Circles highlight areas of cytoplasm with and without bright SG foci. Scale bar = 50 μm. (L) Fraction of transfected cells with silvestrol-induced SGs quantified from K (based on G3BP2 staining). Each data point represents independent biological replicate (N=3). Error bars = standard deviation. One-way ANOVA and Tukey multiple comparisons tests were done to determine statistical significance (***, p -value < 0.001; **, p-value <0.01; ns, non-significant).

Alternatively, using TIAR or G3BP1 as SG markers in these cells could compromise visualization of SGs that may lack these proteins. To distinguish between these possibilities, we stained for G3BP2, another well established SG marker. Unlike with G3BP1 staining, we did not see significant decrease in G3BP2 signal in either OC43 or CoV2 Nsp1 expressing cells, instead we observed that some CoV2 Nsp1 expressing cells were forming small G3BP2-positive SGs upon As treatment (Fig. 5E). We analysed the number and size of As-induced G3BP2-positive SGs that form in CoV2 Nsp1 expressing cells and compared them to SGs formed in control EGFP expressing cells. This analysis revealed that SGs that do form in many CoV2 Nsp1-expressing cells are significantly smaller than SGs that form in control cells, while their average number remains the same (Fig. 5F).

We observed depletion of G3BP1 protein and mRNA in CoV2 infected cells. Next, we tested if G3BP1 depletion by CoV2 Nsp1 but not by OC43 Nsp1 is linked to mRNA degradation induced by the former. We generated two CoV2 Nsp1 amino acid substitution mutants that are defective for mRNA degradation function but are still able to inhibit host protein synthesis: R99A N-terminal domain mutant (99A) and R124A,K125A linker region double mutant (125A) (42). We transiently transfected 293A cells with the OC43 Nsp1, the wild type CoV2 Nsp1, with mutant CoV2 Nsp1 constructs, or with EGFP control and induced SG formation in these cells 24 h post-transfection using As treatment. Immunofluorescence microscopy analysis revealed that all Nsp1 constructs were able to inhibit As-induced SG formation to various degrees (Fig.5G). However, only the wild type CoV2 Nsp1 caused a dramatic increase in nuclear TIAR signal, while OC43 Nsp1, CoV2 99A, and 125A mutants did not affect subcellular TIAR distribution compared to EGFP control (Fig. 5G). Western blot analysis of whole cell lysates revealed that none of the Nsp1 constructs altered total TIAR protein levels, indicating that like CoV2 infection, the wild type CoV2 Nsp1 expression alters nucleocytoplasmic shuttling of TIAR without affecting its expression (Fig. 5H). In addition, only the wild type CoV2 Nsp1 overexpression caused depletion of G3BP1 protein (Fig. 5H, lane 3). This alteration in G3BP1 levels was not observed in OC43 Nsp1- expressing cells or cells expressing CoV2 Nsp1 mutants defective in stimulating mRNA degradation (Fig. 5H, lanes 2,4,5). These results strongly link CoV2 Nsp1-mediated host mRNA degradation to both the nuclear accumulation of TIAR and the depletion of G3BP1 protein. Indeed, when we analyzed host transcript levels in cells transfected with Nsp1 constructs by RT-QPCR, we confirmed that only the wild type CoV2 Nsp1 caused decrease in G3BP1, G3BP2, and β-actin (ACTB) mRNAs (Fig. 5J). This indicated that unlike CoV2 Nsp1, the OC43 Nsp1 does not cause mRNA degradation, and that the amino acid substitution mutants we generated behave as expected. Interestingly, none of the Nsp1 constructs decreased TIAR transcript levels, with the wild type CoV2 Nsp1 causing modest but statistically significant increase in TIAR mRNA compared to EGFP control (Fig. 5J).

To measure and compare SG inhibition by our panel of Nsp1 constructs, we quantified As-induced SG formation using different markers. We saw that when SGs were stained using TIAR as a marker, like in Fig. 5A and 5G, the wild type CoV2 Nsp1 appeared significantly better at preventing SG formation compared to 99A or 125A mutants that did not cause nuclear accumulation of TIAR (Fig. 5I). By contrast, when we quantified G3BP2-positive SGs, wild type and mutant Nsp1 constructs inhibited SG formation to the same degree (Fig. 5F). While all Nsp1 constructs inhibited SG formation visualized using G3BP2 as a marker, more than 50% of cells still formed SGs. Therefore, it is apparent that in many wild type CoV2 Nsp1-expressing cells SGs still form, but they contain very little TIAR and G3BP1.

To examine if inhibition of eIF2α phosphorylation is the main mechanism of SG suppression by OC43 Nsp1, we treated OC43 Nsp1, CoV2 Nsp1, and EGFP expressing cells with Silvestrol. To visualize SGs we used G3BP2 and TIA-1 markers. Bright SG foci formed in the cytoplasm of EGFP-expressing and untransfected bystander cells, as well as in cells expressing OC43 Nsp1 (Fig. 5K). At the same time, most CoV2 Nsp1 expressing cells had either smaller dispersed SG foci or no discernable SGs (Fig. 5K). We quantified the fraction of cells with Silvestrol-induced SGs in all three conditions and demonstrated that only CoV2 Nsp1 decreased SG formation (Fig. 5L). This indicates that mRNA and G3BP1 depletion, as well as nuclear retention of TIAR by CoV2 Nsp1 contribute to impaired SG condensation independent of eIF2α phosphorylation inhibition, with roughly 25% of transfected cells not forming SGs and the remaining cells forming smaller SGs. OC43 Nsp1, on the other hand, did not significantly inhibit SG formation when induced by a mechanism that is independent of eIF2α phosphorylation. Taken together, our experiments show that both OC43 and CoV2 Nsp1 inhibit As-induced SG formation. They both act upstream by inhibiting eIF2α phosphorylation, and in the case of CoV2 Nsp1, downstream by affecting SG nucleation and composition. The mRNA degradation stimulated by CoV2 Nsp1 that causes nuclear accumulation of TIAR and depletion of G3BP1 protein prevents efficient SG condensation, leading to formation of smaller granules lacking TIAR and G3BP1.

### G3BP1 overexpression interferes with OC43 infection

G3BP1 is one of the most important SG nucleating proteins. Apart from its function in nucleating SG formation it is also involved in antiviral signaling. Since our work revealed that coronaviruses are efficient at blocking SG condensation and that CoV2 host shutoff causes G3BP1 depletion, we decided to test if G3BP1 overexpression would be detrimental to virus replication. We generated 293A cells stably transduced with lentivirus vectors encoding EGFP-tagged G3BP1 (293A[EGFP-G3BP1]) and control cells expressing EGFP (239A[EGFP]). To ensure similar levels of expression of these constructs, we sorted early passage cells to have both cell lines with similar GFP signal intensity. Although transient overexpression of G3BP1 may trigger SG formation in the absence of exogenous stress (55), we previously confirmed that this approach generated stable cell lines that did not form SGs spontaneously (30, 56). Initial testing of these cell lines revealed no major differences in cell morphology or SG formation following As treatment (Fig.6A). We infected 293A[EGFP], 293A[EGFP-G3BP1], and parental untransduced 293A cells with the same OC43 virus inoculum at the MOI = 1.0 and analyzed infection rates using immunofluorescence microscopy staining at 24 hpi. This analysis revealed that 293A[EGFP-G3BP1] cells were significantly more resistant to virus infection then either parental or control 293A[EGFP] cells (Fig. 6B,C). Importantly, this was not due to lentiviral integration or non-specific effect of EGFP overexpression as infection rates were the same between 293A[EGFP] and parental untransduced cells (Fig. 6B,C). Western blot analysis confirmed that the EGFP-G3BP1 fusion protein was expressed at higher levels than the endogenous G3BP1 (Fig. 6D). The ectopic overexpression of EGFP-G3BP1 but not the EGFP control caused noticeable decrease in endogenous G3BP1 and G3BP2 expression, but the total level of G3BP1 still remained much higher. Consistent with lower infection rates observed in 293A[EGFP-G3BP1] cells, the accumulation of viral N protein was decreased as well (Fig. 6D). To examine whether the increased resistance of 293[EGFP-G3BP1] cells to coronavirus infection was due to increased SG formation, we analyzed infected cells at 24 hpi using immunofluorescence microscopy with staining for G3BP2 in addition to immunostaining for viral N protein. We did not observe increased SG formation in 293[EGFP-G3BP1] cells compared to either control, with most cells remaining SG-free (Fig. 6E). This result suggests that OC43 virus is effective at suppressing SG formation even when G3BP1 levels are elevated. This also indicates that the antiviral function of G3BP1 is not limited to SG nucleation.

**Figure 6.**
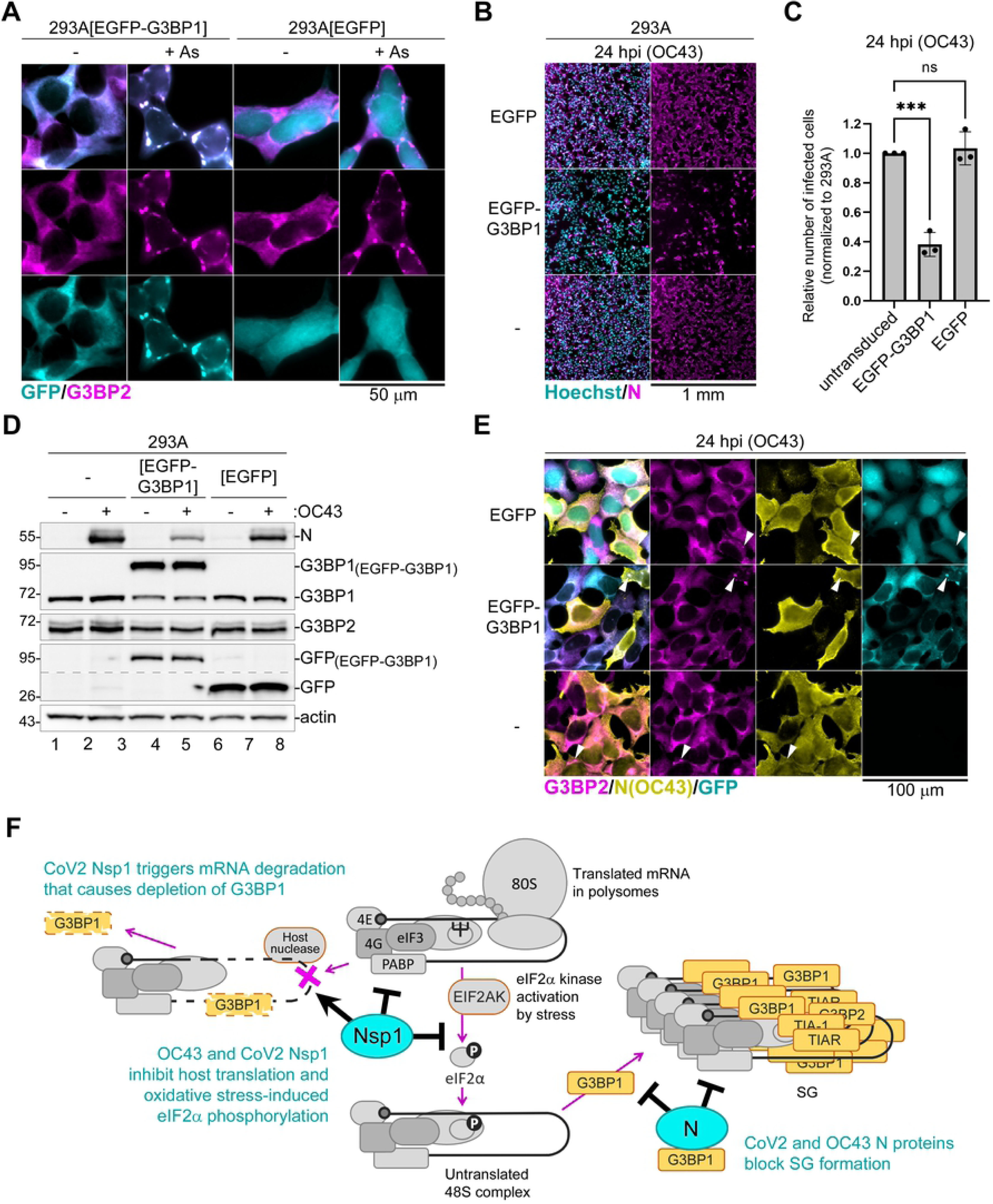
G3BP1 overexpression inhibits OC43 infection independent from SG formation. (A) Immunofluorescence microscopy analysis of 293A[EGFP] and 293A[EGFP-G3BP1] cells untreated (-) or treated with arsenite (+ As) and stained for G3BP2 (magenta). GFP signal is shown in teal. Scale bar = 50 µm. (B) Immunofluorescence microscopy analysis of parental 293A cells (-), 293A[EGFP], and 293A[EGFP-G3BP1] cells infected with OC43 at MOI = 1.0 and stained for OC43 N protein (magenta) at 24 hpi. Nuclei were stained with Hoechst dye (teal). Scale bar = 1 mm. (C) Relative number of infected cells (normalized to 293A) in 293A[EGFP] and 293A[EGFP-G3BP1] cells at 24 hpi, quantified from E. Each data point represents independent biological replicate (N=3). Error bars = standard deviation. One-way ANOVA and Dunnett’s multiple comparisons tests were done to determine statistical significance (***, p -value < 0.001, ns, non-significant). (D) Western blot analysis of 293A, 293A[EGFP], and 293A[EGFP-G3BP1] cells infected with OC43 at 24 hpi. Anti-GFP blot was spliced along the dotted line to reduce vertical size. (E) Immunofluorescence microscopy of cells infected as in B and immunostained for G3BP2 (magenta) and OC43 N (yellow). GFP signal is shown in teal. Arrowheads indicate infected cells that formed SGs. Scale bar = 100 µm. (F) Diagram illustrating concerted action of N and Nsp1 proteins in disrupting G3BP1 activity and inhibiting SG formation. Poly(A) binding protein (PABP) and the eukaryotic translation initiation factors eIF3, eIF4E (4E), and eIF4G (4G) that form preinitiation complex and recruit ribosomal subunits are shown schematically. EIF2AK = eIF2α kinase.

## Discussion

Despite high prevalence in the population and ability to cause reinfections (57, 58), common cold human coronaviruses remained poorly studied due to lower clinical significance compared to other seasonal respiratory viruses like influenza or respiratory syncytial virus (RSV). With the emergence of highly pathogenic coronaviruses of zoonotic origin, especially the recent pandemic CoV2 that swept the globe causing high morbidity and mortality, the interest in coronavirus research increased. Due to its classification as level 2 pathogen, OC43 emerged as one of the model coronaviruses (59–63). In this study, we examined this viruses’ ability to inhibit formation of SGs, one of the intrinsic host antiviral mechanisms, and compared it to that of the highly pathogenic CoV2.

Our work demonstrates that in virus-infected cells SGs do not form until about 24 h post-infection, with fewer than 5% of cells having SGs at that time point. Despite coronaviruses being known to generate levels of dsRNA sufficient to be detected by dsRNA-specific antibody (17, 64), both OC43 and CoV2 do not activate PKR to levels that would induce significant eIF2α phosphorylation and SG formation.

Furthermore, both viruses inhibit eIF2α phosphorylation triggered by As treatment which causes oxidative stress and activates HRI (65). This indicates that OC43 and CoV2 suppress eIF2α phosphorylation-dependent translation arrest. The nuclease activity of Nsp15, conserved in coronaviruses from different genera, was previously shown to be important in limiting dsRNA detection in cells infected with Gammacoronavirus IBV (17). Infection of cells with a recombinant virus with H238A substitution in Nsp15 that abrogates its nuclease activity triggered PKR activation, phosphorylation of eIF2α, and induction of SGs (17). Thus, it is very likely that Nsp15 activity also contributes to the lack of PKR activation and SG formation in OC43 and CoV2 infected cells. Despite this, in our experimental system, overexpression of OC43 Nsp15 did not significantly affect As-induced eIF2α phosphorylation or SG formation. In another study, CoV2 N and the 3CLpro proteinase Nsp5 were shown to suppress SGs induced by transfection with the dsRNA mimic polyinosinic-polycytidylic acid (poly(I:C)) (34).

However, the effects of these proteins on poly(I:C)-induced PKR activation and eIF2α phosphorylation were not examined. Previous studies have demonstrated that CoV2 N protein directly binds G3BP1 and interferes with its function in SG nucleation (33, 34). Consistent with this mechanism, upon ectopic overexpression, CoV2 N protein blocked both phospho-eIF2α dependent and independent SG formation in our study. In comparison, the OC43 N protein was also able to inhibit SGs but was better at inhibiting phospho-eIF2α independent SGs induced by Sil. than the SGs induced by As. Despite these differences in magnitude, our results show that inhibition of SG nucleation by N protein is conserved between OC43 and CoV2. Notably, neither N protein affected As-induced eIF2α phosphorylation, confirming that they function downstream at the SG nucleation step. Instead, our work revealed that OC43 and CoV2 Nsp1 host shutoff factors are responsible, at least in part, for inhibition of eIF2α phosphorylation observed in As-treated infected cells. Upon ectopic overexpression, OC43 and CoV2 Nsp1 proteins inhibited eIF2α phosphorylation and SG formation induced by As. Importantly, OC43 Nsp1 did not significantly inhibit Sil.-induced SG formation, indicating that inhibition of p-eIF2α mediated translation arrest is the main mechanism of SG inhibition by this protein. By contrast, CoV2 Nsp1 was also able to inhibit Sil.-induced SG formation, although to a much lesser extent compared to As-induced SGs. Since virus infections or CoV2 or OC43 Nsp1 overexpression did not affect total eIF2α levels in our experiments, suppression of eIF2α phosphorylation by Nsp1 could be through direct inhibition of a specific kinase (e.g. HRI) or through stimulation of eIF2α dephosphorylation. It is important to note, however, that our results do not rule out the contribution of other viral proteins to suppression of eIF2α phosphorylation in infected cells.

Another striking phenotype that we observed in CoV2 but not OC43 infected cells was the depletion of G3BP1 protein and increase in nuclear retention of TIAR. Our analysis using translation inhibitors revealed that G3BP1 protein is not intrinsically unstable, indicating that general inhibition of protein synthesis by CoV2 host shutoff is not responsible for G3BP1 depletion. Instead, we discovered that G3BP1 levels can be decreased by transcription inhibitor ActD. This suggests a link between general cytoplasmic mRNA depletion by CoV2 host shutoff factor Nsp1 and the sharp decrease in G3BP1 protein levels. Indeed, upon ectopic overexpression, CoV2 Nsp1 caused depletion of G3BP1 protein level. By contrast, OC43 Nsp1, which does not induce mRNA degradation, did not affect G3BP1 expression. To firmly link G3BP1 protein depletion and nuclear TIAR accumulation to mRNA degradation induced by CoV2 Nsp1, we created and tested two amino acid substitution mutants that were previously shown to be defective in stimulating host mRNA degradation – single amino acid substitution R99A in the Nsp1 N-terminal domain, and double substitution R124A,K125A in the linker region (42). Neither mutant caused decrease in host mRNA levels nor affected G3BP1 protein levels or nuclear TIAR accumulation, similar to OC43 Nsp1.

G3BP1 protein and its homologue G3BP2 are master regulators of SG formation that can directly interact with the small ribosomal subunit and facilitate initial nucleation of SGs (26,55,66). Consequently, most types of stress fail to induce SG formation in G3BP1/G3BP2 double knock-out cells (26, 66). Interestingly, despite partially overlapping functions, silencing of either G3BP1 or G3BP2 can inhibit both phospho-eIF2α dependent and independent SG formation, suggesting that the total levels of G3BP1/G3BP2 expression affect SG nucleation (26,56,67). In addition, G3BP1 protein is involved in antiviral responses through multiple mechanisms (29,68–70). It can amplify translation arrest by recruiting unphosphorylated PKR to stress granules, where it becomes activated in a dsRNA-independent manner (29). It is also involved in the activation of signaling cascades leading to induction of antiviral cytokines (29, 70), and many studies have shown that silencing of G3BP1 leads to impaired induction of type I interferon (27,29,71). Recently, it was discovered that G3BP1 and G3BP2 function in anchoring the tuberous sclerosis complex (TSC) to lysosomes and suppressing activation of the mechanistic target of rapamycin complex 1 (mTORC1) (72). These functions of G3BP1 are independent from SG formation. Thus, in addition to interfering with SG nucleation, depletion of G3BP1 in CoV2 infected cells could benefit viral replication by both blunting the cellular innate immune responses and by upregulating biosynthetic pathways through relieving mTORC1 suppression. In our study we showed that overexpression of G3BP1 inhibits OC43 infection without increasing SG formation, suggesting that G3BP1 is antiviral towards OC43 and that some of the SG nucleation-independent functions of G3BP1 could be responsible.

Existence of multiple mechanisms that interfere with translation arrest and SG formation in cells infected with both the seasonal common cold OC43 and the pandemic CoV2 viruses described in this study highlights the importance of overcoming these antiviral mechanisms by diverse coronaviruses. In addition, our work discovers a novel feature of Nsp1-mediated host shutoff that simultaneously blocks host translation initiation and promotes continuous regeneration of GTP-bound translation initiation-competent eIF2 by inhibiting eIF2α phosphorylation. In the follow up work focusing on CoV2 and OC43 Nsp1, we aim at characterizing the mechanism by which these host shutoff factors inhibit eIF2α phosphorylation and the contribution of this function to viral replication fitness and suppression of host antiviral responses.

## Materials and Methods

### Cells

Human Embryonic Kidney (HEK) 293A cells and human colon adenocarcinoma (HCT-8) cells were cultured in Dulbecco’s modified Eagle’s medium (DMEM) supplemented with heat-inactivated 10% fetal bovine serum (FBS), and 2 mM L-glutamine (all purchased from Thermo Fisher Scientific (Thermo), Waltham, MA, USA). BEAS-2B cells were cultured in Bronchial Epithelial Cell Growth Medium (BEGM, Lonza, Kingston, ON, Canada) on plates prepared with coating media (0.01 mg/ml fibronectin, 0.03 mg/mL bovine collagen type I, and 0.01 mg/ml bovine serum albumin (all from Millipore Sigma, Oakville, ON, Canada) dissolved in Basal Epithelial Cell Growth Medium (BEBM, Lonza)). 293A and BEAS-2B cells were purchased from American Type Culture Collection (ATCC, Manassas, VA, USA), HCT-8 cells were purchased from Millipore Sigma.

### Viruses

HCoV-OC43 was purchased from ATCC and SARS-CoV2 (strain SARS-CoV-2/SB3-TYAGNC) was derived from a clinical isolate and generously provided by Drs. Arinjay Banerjee, Karen Mossman and Samira Mubareka (73). To generate initial HCoV-OC43 virus stocks, Vero E6 cells (ATCC) were infected at multiplicity of infection (MOI) <0.1 for 1 h in serum-free DMEM at 37°C following replacement of the inoculum with DMEM supplemented with 1% FBS and continued incubation at 33°C. Once CPE reached 75% at 4-5 d past-infection, the viral supernatant was harvested, centrifuged at 2,500 x g for 5 min, and then the cleared viral supernatant was aliquoted and stored at -80°C. For SARS-CoV-2 stocks, Vero E6 cells in a confluent T-175 cm2 flask were infected at a MOI of 0.01 for 1 h at 37°C in 2.5 mL of serum-free DMEM with intermittent shaking every 10 min. Following incubation, 17.5 ml of DMEM supplemented with 2% FBS was added directly to the viral inoculum and continued incubation at 37°C. With the onset of CPE at 62 – 66 hpi, viral supernatant was harvested, centrifuged at 1,000 x g for 5 min, and then the cleared viral supernatant was aliquoted and stored at -80°C. Stocks were titered by plaque assay on Vero E6 cells as in (74).

### Plasmids and lentivirus stocks

SARS-CoV2 and HCoV-OC43 N, Nsp1, and Nsp15 open reading frames were PCR-amplified from cDNAs generated from total RNA of infected Vero E6 cells collected at 24 hpi using specific primers with simultaneous introduction of flanking restriction sites. Then, coding sequences were inserted between EcoRI and XhoI sites into pCR3.1-EGFP vector (75) to generate pCR3.1-EGFP-OC43-N, pCR3.1-EGFP-CoV2-N, pCR3.1-EGFP-OC43-Nsp1, pCR3.1-EGFP-CoV2-Nsp1, and pCR3.1-EGFP-OC43-Nsp15 plasmids. To generate N-terminally HA-tagged Nsp1 constructs, coding sequences were inserted between KpnI and XhoI sites into 3xHA-miniTurbo-NLS_pCDNA3 vector (a gift from Alice Ting, Addgene plasmid # 107172) to generate pCDNA3-HA-OC43-Nsp1 and pCDNA3-HA-CoV2-Nsp1 vectors (miniTurbo-NLS coding sequence was replaced by Nsp1 sequences). Amino acid substitutions in pCDNA3-HA-CoV2-Nsp1 vector were introduced using Phusion PCR mutagenesis (New England Biolabs) to generate pCDNA-HA-CoV2-Nsp1(R99A) and pCDNA-HA-CoV2-Nsp1(R124A,K125A) vectors. To generate lentivirus vectors pLJM1-ACE2-BSD, pLJM1-EGFP-BSD, and pLJM1-EGFP- G3BP1-BSD, the PCR-amplified ACE2, EGFP, and G3BP1 coding sequences were inserted into the multicloning site of pLJM1-B* vector (76). All constructs were verified by Sanger sequencing, sequences are available upon request. To generate lentivirus stocks, HEK 293T cells (ATCC) were reverse-transfected with polyethylenimine (PEI, Polysciences, Warrington, PA, USA) and the following plasmids for lentiviral generation: pLJM1-B* backbone-based constructs, pMD2.G, and psPAX2. pMD2.G and psPAX2 are gifts from Didier Trono (Addgene plasmids #12259 and #12260). 48 h post-transfection, lentivirus containing supernatants were passed through a 0.45 μm filter and frozen at -80⁰C.

### Generation of stably transduced cell lines

To generate 293A-ACE2 cells, 293A cells were stably transduced with a lentivirus vector encoding ACE2 (pLJM1-ACE2-BSD) and selected and maintained in 10 μg/mL Blasticidin S HCl (Thermo Fisher). To generate 293A[EGFP] and 293A[EGFP-G3BP1] cells, 293A cells were transduced with lentiviruses produced from pLJM1-EGFP-BSD and pLJM1-EGFP-G3BP1-BSD vectors and at passage 3 post-transduction, EGFP-positive cells were isolated using live cell sorting on BD FACSAria III instrument, cultured and used for experiments at passage 5 to 6.

### Cell treatments

For SG induction, sodium arsenite (Millipore Sigma) was added to the media to a final concentration of 500 µM and cells were returned to 37⁰C incubator for 50 min; Silvestrol (MedChemExpress, Monmouth Junction, NJ, USA) was added to the media to a final concentration of 500 nM and cells were returned to 37⁰C incubator for 1 h. For treatment of 293A cells with translation and transcription inhibitors, cycloheximide (50 µg/ml), Actinomycin D (5 µg/ml), or Silvestrol (320 nM) were added to the media and cells were incubated for 12h prior to lysis for western blot.

### Virus infections

Cell monolayers were grown in 20-mm wells of 12-well cluster dishes with or without glass coverslips. For HCoV-OC43 infections, media was aspirated, cells were washed briefly with PBS and 300 µl of virus inoculum diluted to the calculated MOI = 1.0 in 1% FBS DMEM was added. Cells were placed at 37°C for 1 h, with manual horizontal shaking every 10-15 minutes. Then, virus inoculum was aspirated from cells, cells were washed with PBS, 1 ml of fresh 1% FBS DMEM was added to each well, and cells were returned to 37°C until the specified time post-infection. For SARS-CoV-2 infections, media was aspirated and 100 μL of virus inoculum diluted in serum-free DMEM at a MOI of 0.2 was added directed to the wells. Cells were incubated at 37°C for 1h with intermittent shaking every 10 min. Following incubation, virus inoculum was removed, and cells were washed with 1 mL of PBS three times, then 1 mL of fresh DMEM supplemented with 10% FBS was added to each well. Cells were incubated at 37°C for 24 h.

### Transfection

293A cells were seeded into 20-mm wells of 12-well cluster dishes with or without glass coverslips and the next day transfected with 500 ng DNA mixes/well containing expression vectors (250 ng) and pUC19 filler DNA (250 ng) using Fugene HD (Promega, Madison, WI, USA) according to manufacturer’s protocol. Where indicated, the amount of filler DNA was reduced to 150 ng and 100 ng of the pCR3.1- EGFP plasmid was co-transfected with expression vectors for Nsp1 proteins. Cells were used for experiments 23-24 h post-transfection as indicated.

### Immunofluorescence staining

Cell fixation and immunofluorescence staining were performed according to the procedure described in (56). Briefly, cells grown on 18-mm round coverslips were fixed with 4% paraformaldehyde in PBS for 15 min at ambient temperature and permeabilized with cold methanol for 10 min. After 1-h blocking with 5% bovine serum albumin (BSA, BioShop, Burlington, ON, Canada) in PBS, staining was performed overnight at +4°C with antibodies to the following targets: CoV2 N (1:400; rabbit, Novus Biologicals, NBP3-05730); eIF3B (1:400; rabbit, Bethyl Labs, A301761A); eIF4G (1:200; rabbit, Cell Signaling,

#2498); G3BP1 (1:400; mouse, BD Transduction, 611126); G3BP2 (1:1000; rabbit, Millipore Sigma, HPA018304); HA tag (1:100; mouse, Cell Signalling, #2367); OC43 N (1:500; mouse, Millipore, MAB9012); TIA-1 (1:200; goat, Santa Cruz Biotechnology, sc-1751); TIAR (1:1000; rabbit, Cell Signaling, #8509). Alexa Fluor (AF)-conjugated secondary antibodies used were: donkey anti-mouse IgG AF488 (Invitrogen, A21202), donkey anti-rabbit IgG AF555 (Invitrogen, A31572), donkey anti-goat IgG AF647 (Invitrogen, A32839). Where indicated, nuclei were stained with Hoechst 33342 dye (Invitrogen, H3570). Slides were mounted with ProLong Gold Antifade Mountant (Thermo Fisher) and imaged using Zeiss AxioImager Z2 fluorescence microscope and Zeiss ZEN 2011 software. Green, red, blue, and far-red channel colors were changed for image presentation in the color-blind safe palette without altering signal levels. Quantification of SG-positive cells was performed by counting the number of cells with at least two discrete cytoplasmic foci from at least 3 randomly selected fields of view, analysing >100 cells per treatment in each replicate. Analysis of SG number and size was performed on cropped images of individual cells using ImageJ software Analyze Particles function after automatic background substraction and thresholding. For each of 3 independent biological replicates, 7 cells selected from at least 3 random fields of view were analyzed for a total of 21 cells per condition.

### Western blotting

Whole-cell lysates were prepared by direct lysis of PBS-washed cell monolayers with 1× Laemmli sample buffer (50 mM Tris-HCl pH 6.8, 10% glycerol, 2% SDS, 100 mM DTT, 0.005% Bromophenol Blue). Lysates were immediately placed on ice, homogenized by passing through a 21-gauge needle, and stored at −20°C. Aliquots of lysates thawed on ice were incubated at 95°C for 3 min, cooled on ice, separated using denaturing PAGE, transferred onto PVDF membranes using Trans Blot Turbo Transfer System with RTA Transfer Packs (Bio-Rad Laboratories, Hercules, CA, USA) according to manufacturer’s protocol and analysed by immunoblotting using antibody-specific protocols. Antibodies to the following targets were used: β-actin (1:2000; HRP-conjugated, mouse, Santa Cruz Biotechnology, sc- 47778); CoV2 N (1:1,000; rabbit, Novus Biologicals, NBP3-05730); eIF2α (1:1000; rabbit, Cell Signaling, #5324); eIF4G (1:1000; rabbit, Cell Signaling, #2498); G3BP1 (1:4000; mouse, BD Transduction, 611126); G3BP2 (1:2500; rabbit, Millipore Sigma, HPA018304); GFP (1:1000; rabbit, Cell Signaling, #2956); HA tag (1:1,000; mouse, Cell Signalling, #2367); OC43 N (1:1,000; mouse, Millipore, MAB9012); phospho-S51-eIF2α (1:1000; rabbit, Cell Signaling, #3398); TIAR (1:1000; rabbit, Cell Signaling, #8509). For band visualization, HRP-conjugated anti-rabbit IgG (Goat, Cell Signaling, #7074) or anti-mouse IgG (Horse, Cell Signaling, #7076) were used with Clarity Western ECL Substrate on the ChemiDoc Touch Imaging Sysytem (Bio-Rad Laboratories). Where indicated, total protein was visualised post-transfer to PVDF membranes on ChemiDoc using Stain-free fluorescent dye (Bio-Rad Laboratories). For analyses of protein band intensities, western blot signals were quantified using Bio-Rad Image Lab 5.2.1 software and values normalized to the Stain-free signal for each lane.

### Ribopuromycylation assay

The puromycin incorporation assay was performed as described in (77) with the following modifications. Puromycin was added to the medium at the final concentration of 10 μg/ml for 10 min. Cells were washed with PBS and the whole-cell lysate preparation and western blotting analysis were done as described above. For electrophoresis, samples were loaded onto Mini-PROTEAN TGX Pre-cast Stain-Free gels (5-15%, BioRad Laboratories, Hercules, CA, USA) and total protein was visualised post-transfer to PVDF membranes on ChemiDoc Touch Imaging System. Puromycin incorporation into nascent polypeptides was visualised using anti-puromycin antibody (1:6,000; mouse, MilliporeSigma, MABE343).

### RNA isolation and RT-QPCR

Total RNA was isolated from cells using RNeasy Plus Mini kit (Qiagen) according to manufacturer’s protocol. 250 ng of RNA was used to synthesize cDNA using qScript cDNA SuperMix (Quanta) or Maxima H Minus Reverse Transcriptase (Thermo Fisher). Quantitative PCR amplification was performed using PerfeCTa SYBR Green PCR master mix (Quanta) and specific primers listed below on Cielo 3 QPCR unit (Azure). Primers used: 18S - Left: cgttcttagttggtggagcg, Right: ccggacatctaagggcatca; ACTB - Left: catccgcaaagacctgtacg, Right: cctgcttgctgatccacatc; G3BP1 - Left: ggtcttaggcgtgtaccctg, Right: tatcgggaggaccctcagtg; G3BP2 - Left: gcctgttaatgctgggaacac, Right: tgttgcctcctgttgcagat; TIAR - Left: tggaagatgcagaagaccgag, Right: tgcactccctagctctgaca. Relative target levels were determined using ΔΔCt method with normalization to 18S.

### Statistical analyses

All numerical values are plotted as means (bar graphs) and display individual datapoints representing independent biological replicates (separate experiments performed on different days); the error bars represent standard deviations. Statistical analyses for each data set are described in figure legends and were performed using GraphPad Prism 8 software.

## Acknowledgements

We would like to thank Dr. Arinjay Banerjee (Vaccine and Infectious Disease Organization (VIDO), Dr. Karen Mossman (University of McMaster) and Dr. Samira Mubareka (University of Toronto) for the SARS-CoV-2 isolate. We also thank Dalhousie CORES Flow Cytometry Facility for assistance in generating stable cell lines expressing EGFP-G3BP1 and EGFP. Finally, we thank members of Khaperskyy and Corcoran labs for helpful discussions about experimental design and their critical input on the draft manuscript. This work was supported by Canadian Institutes for Health Research (CIHR) Project Grant PJT-175130 and Research Nova Scotia Grant RNS-NHIG-2020-1383 (to D.K.). This study was also supported in part by operating funds awarded to JAC from the Canadian Institutes for Health Research (CIHR): a COVID rapid response operating grant (#177704) and an operating grant (#175622) the Coronavirus Variants Rapid Response Network (CoVaRR-Net), of which JAC is a member. MK was supported by a CSM graduate training award and a CIHR doctoral award.

## Notes

### Competing Interest Statement

The authors have declared no competing interest.

